# Time of day is associated with paradoxical reductions in global signal fluctuation and functional connectivity

**DOI:** 10.1101/653899

**Authors:** Csaba Orban, Ru Kong, Jingwei Li, Michael W.L. Chee, B. T. Thomas Yeo

**Affiliations:** Department of Electrical and Computer Engineering, ASTAR-NUS Clinical Imaging Research Centre, Singapore Institute for Neurotechnology and Memory Networks Program, National University of Singapore, Singapore; Neuropsychopharmacology Unit, Centre for Psychiatry, Imperial College London, London, UK; Centre for Cognitive Neuroscience, Duke-NUS Medical School, Singapore; Martinos Center for Biomedical Imaging, Massachusetts General Hospital, Charlestown, MA, USA; NUS Graduate School for Integrative Sciences and Engineering, National University of Singapore, Singapore

**Keywords:** time of day, circadian, diurnal, global signal, fMRI, resting state, human connectome project

## Abstract

The brain exhibits substantial diurnal variation in physiology and function but neuroscience studies rarely report or consider the effects of time of day. Here, we examined variation in resting-state fMRI in around 900 subjects scanned between 8am to 10pm on two different days. Multiple studies across animals and humans have demonstrated that the brain’s global signal amplitude (henceforth referred to as “fluctuation”) increases with decreased arousal. Thus, in accord with known circadian variation in arousal, we hypothesised that global signal fluctuation would be lowest in the morning, increase in the mid-afternoon and dip in the early evening. Instead, we observed a cumulative decrease (22% between 9am to 9pm) in global signal fluctuation as the day progressed. To put the magnitude of this decrease in context, we note that task-evoked fMRI responses are typically in the order of 1% to 3%. Respiratory variation also decreased with time of day, although control analyses suggested that this did not account for the reduction in GS fluctuation. Finally, time of day was associated with marked decreases in resting state functional connectivity across the whole brain. The magnitude of decrease was significantly stronger than associations between functional connectivity and behaviour (e.g., fluid intelligence). These findings reveal unexpected effects of time of day on the resting human brain, which challenge the prevailing notion that the brain’s global signal reflects mostly arousal and physiological artefacts. We conclude by discussing potential mechanisms for the observed diurnal variation in resting brain activity and the importance of accounting for time of day in future studies.

## 2. Introduction

Circadian rhythms govern diverse aspects of physiology including sleep/wake cycles (Dijk and Lockley, 2002), cognition (Schmidt et al., 2007), gene expression (Storch et al., 2002), temperature regulation (Refinetti and Menaker, 1992) and endocrine signalling (Asher and Schibler, 2011). Similarly, studies of brain function in both humans and animals have documented time of day-dependent variation at multiple scales of brain organization. Examples include diurnal variation in synaptic protein levels and synaptic strength (Vyazovskiy et al., 2008; Gilestro et al.,2009; Liu et al., 2010), the amplitude and slope of cortical or motor evoked responses (Vyazovskiy et al., 2008; Huber et al., 2013; Kuhn et al., 2016), theta power (Finelli et al., 2000; Hung et al., 2013) as well as changes in cerebral blood flow (Braun, 1997; Hodkinson et al., 2014; Elvsåshagen et al., 2019), task-related BOLD activation (Schmidt et al., 2009; Muto et al., 2016; Byrne et al., 2017), resting state BOLD signal amplitude (Jiang et al., 2016; Cordani et al., 2018) and resting state functional connectivity (Blautzik et al., 2013; Shannon et al., 2013; Hodkinson et al., 2014).

Despite the clear influence of circadian rhythms on physiology, most studies of brain function do not report or consider the impact of time of day on their findings. Perhaps, there is an implicit assumption in the field that diurnal variation of brain activity is relatively small and unlikely to introduce substantial systematic bias into group analyses. Furthermore, most previous studies of time of day effects have employed small samples, within-subject designs with few time-points, a controlled pre-study routine and in some cases sleep deprivation (e.g. Huber et al., 2013; Muto et al., 2016). Previous resting-state fMRI studies that have focused on time of day have yielded inconsistent results, potentially because of small sample sizes (Blautzik et al., 2013; Shannon et al., 2013; Hodkinson et al., 2014; Jiang et al., 2016; Cordani et al., 2018). Here we exploited resting-state fMRI data from the Human Connectome Project (HCP; Van Essen et al., 2012) involving (i) a large sample of 942 subjects scanned using cutting-edge multiband acquisition sequences, (ii) a wide spread of scan times with a relatively smooth distribution between 9am to 9pm, (iii) test-retest data with varying scan times between two different days (sessions) enabling within-subject analyses, and (iv) individually curated respiratory and cardiac recordings for examination of non-neural effects.

The HCP dataset allowed us to investigate how the magnitude of the brain’s global signal fluctuation (also referred to as amplitude in the literature) varies throughout the day. Our focus on the global signal was motivated by converging evidence of a strong link between global signal fluctuation and arousal/vigilance levels in both humans and non-human primates (reviewed in Liu et al., 2017), which is known to change throughout the day (Van Dongen and Dinges, 2000). Specifically, the magnitude of global signal fluctuation (i.e., its temporal standard deviation) has been found to vary as a function of behavioural and electrophysiological indices of drowsiness (Wong et al., 2013, 2016; Chang et al., 2016) to increase following sleep deprivation (Yeo et al., 2015) and to decrease following caffeine consumption (Wong et al., 2012). Therefore, we hypothesised that levels of global signal fluctuation would be lowest in the late morning hours (e.g. 11am) when arousal levels are normally at their peak (Strogatz et al., 1987; Van Dongen and Dinges, 2000; Bes et al., 2009; de Zeeuw et al., 2018). In addition, we predicted that global signal fluctuation would be elevated during mid-afternoon hours (e.g., 3pm corresponding to the “mid-afternoon dip”) and reduced in the early evening hours (e.g. 8-9pm corresponding to the “wake maintenance zone”), when subjects typically experience reduced and elevated states of arousal respectively (Strogatz et al., 1987; Van Dongen and Dinges, 2000; Bes et al., 2009; de Zeeuw et al., 2018).

The contributions of this study are multi-fold. First, we demonstrated counter-intuitive reductions in the magnitude of global signal fluctuation from late morning to late evening across participants in two separate sessions. Our observation of high levels of global signal fluctuation (typically a feature of low arousal) in the late morning and the gradual reduction in global signal fluctuation throughout the day, were inconsistent with expected patterns of circadian variation in arousal levels. Second, these findings were replicated within-subjects, across two sessions, thus ruling out individual differences in chronotype as a potential explanation. Third, the effect of time of day on global signal fluctuation was not accounted for by differences in head motion, heart rate and respiratory variability. Fourth, the reduction in global signal fluctuation was accompanied by a widespread decrease in inter-regional functional connectivity larger than typical association strength between behaviour (e.g. fluid intelligence) and functional connectivity. In summary, this study revealed robust, diurnal changes in large-scale resting brain activity that merit further investigation.

## 3. Results

### 3.1 Global signal (GS) fluctuation declines with time of day

We considered the MSMAll ICA-FIX fMRI data from HCP S1200 release (Van Essen et al., 2012; Smith et al., 2013). Although there are different definitions of GS in the literature, they all produce highly similar estimates (Carbonell et al., 2011; Smith et al., 2013; Li et al., 2019). Here, we defined the GS to be the mean time-series obtained by averaging the preprocessed fMRI time courses across all cortical grey matter locations (Kong et al., 2019; Li et al., 2019). We also verified that our main findings were robust to using other definitions of GS, e.g., averaging signal across the whole brain. Based on earlier work, we defined GS fluctuation as the temporal standard deviation of the GS (Wong et al., 2012, 2013). We will use the term GS fluctuation instead of the nomenclature of “GS amplitude” from previous studies.

Across participants, GS fluctuation was negatively correlated with time of day for Session 1 (r = - 0.17, p < 1.0e-5, N = 942; Figure 1A) and for Session 2 (r = - 0.20, p < 1.0e-5, N = 869; Figure 1B). While individuals are known to express variation in circadian phase and chronotype (Roenneberg et al., 2007), on average, participants should exhibit higher levels of arousal during late morning (e.g. at 11am) and again in the early evening (e.g. 8-9 p.m.) than in the mid-afternoon (e.g. at 3pm) based on previous studies (Strogatz et al., 1987; Van Dongen and Dinges, 2000; Bes et al., 2009; de Zeeuw et al., 2018). Given the literature suggesting a strong relationship between GS fluctuation and arousal levels (reviewed in Liu et al., 2017), one would expect GS fluctuation to be lower in the late morning (when arousal is generally higher), than in the mid-afternoon (when arousal is generally lower). Instead we found GS fluctuation to exhibit a steady decrease between-subjects from 11am to 9 pm (Figures 1A-B). From 9am to 9pm, GS fluctuation decreased by 22% and 26% in sessions 1 and 2 respectively.

**Figure 1.**
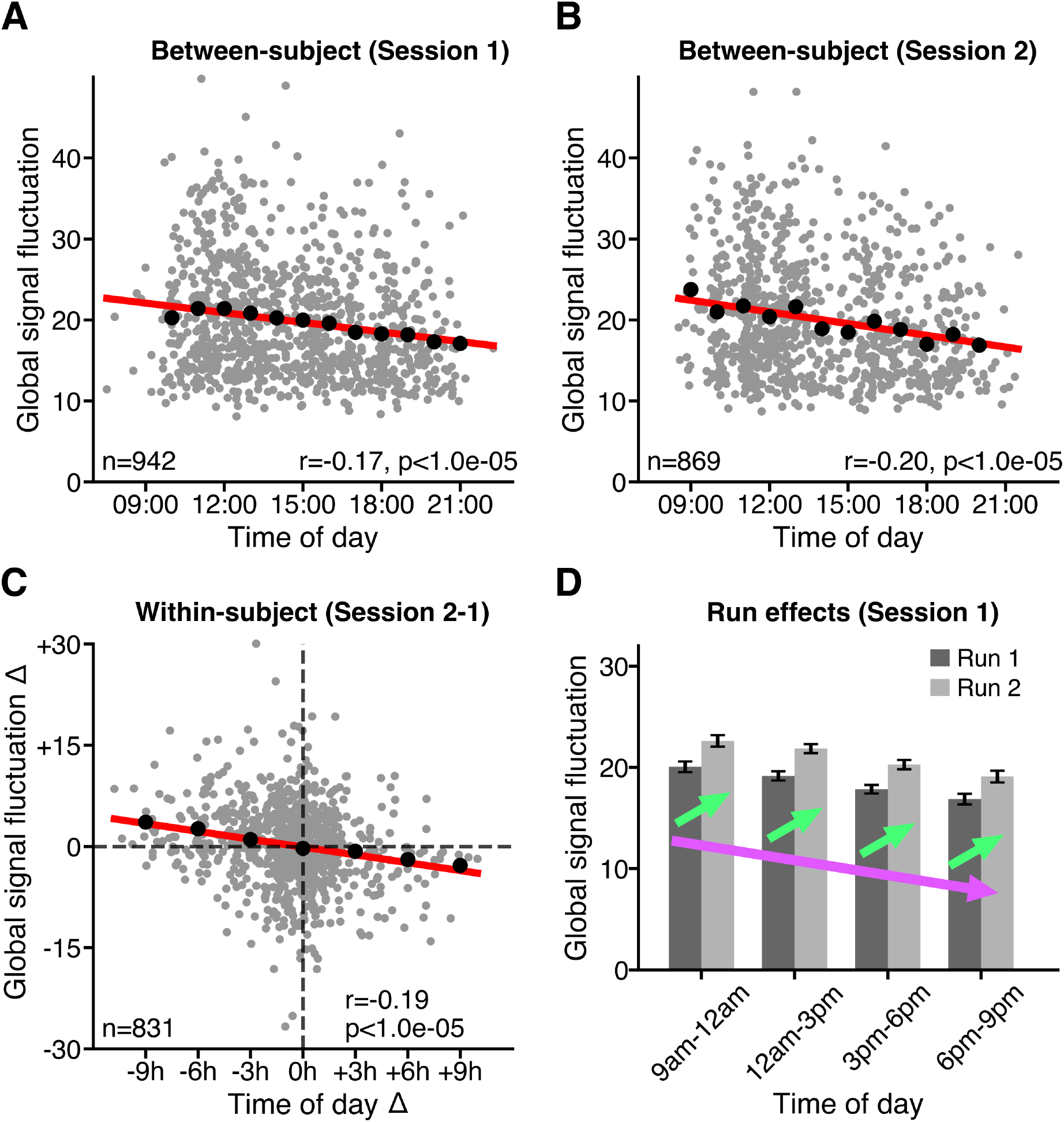
The brain’s global signal (GS) fluctuation (i.e., standard deviation of global signal) decreases with time of day. **(A)** Between-subject variation in GS fluctuation as a function of time of day in Session 1. **(B)** Between-subject variation in GS fluctuation as a function of time of day in Session 2. **(C)** Within-subject variation in GS fluctuation Δ as a function of time of day Δ, where Δ denotes difference between Session 2 and Session 1. Grey dots denote individual subjects. Black dots show mean of GS fluctuation in (A, B) hourly, or (C) 3-hourly time windows. Line of best fit (red) was calculated based on data from all subjects in each plot. R values denote Pearson r correlation coefficient. P values were derived from 100,000 permutations, while keeping family structure intact (Winkler et al., 2015). **(D)** GS fluctuation is elevated in Run 2 compared with Run 1 despite downward shift in GS fluctuation as a function of time of day. Bar plots denote mean GS fluctuation across subjects within 3-hourly time windows for each run. Error bars denote standard error of the mean. Two opposing effects are observable: a fast increase in GS fluctuation on the scale of minutes, i.e. run effect, (green arrows), superimposed on a downward drift of GS fluctuation occurring on the scale of hours, i.e. time of day effect (violet arrow).

To test for within-subject effects, GS fluctuation and time of day from Session 2 were subtracted from their respective values from Session 1 and the resulting values were correlated. Within-subject differences in GS fluctuation as a function of differences in time of day (r = - 0.20, p < 1.0e-5, N = 831), strongly corroborated the between-subject effects (Figure 1C).

Subjects exhibited substantial between-subject variation in GS fluctuation at each time of day. Nevertheless, hourly (black dots in Figure 1A-B) or 3 hourly (black dots in Figure 1C) time-window averages of GS fluctuation closely followed the line of best fit, suggesting a linear decline that was not preferentially driven by any specific time of day.

### 3.2 Scanning run and time of day have opposing effects on GS fluctuation

Within a given session (day), participants were scanned for two consecutive runs. As shown in Figure 1D, GS fluctuation was significantly elevated in Run 2 compared with Run 1 within both Session 1 (t = 13.0, p < 1.0e-05, N = 942) and Session 2 (t = 9.0, p < 1.0e-05, N = 869), consistent with similar analyses of the HCP dataset (Bijsterbosch et al., 2017).

Nevertheless, the negative correlation between time of day and GS fluctuation was significant for each run in both sessions (Figure S1). Thus, in summary, we observed two opposing effects: a fast increase of GS fluctuation occurring on the scale of minutes (i.e. run effect), superimposed on a more gradual decrease of GS fluctuation occurring on the scale of hours (i.e. time of day effect).

### 3.3 Head motion and cardiac measures show strong correlation with GS fluctuation, but not with time of day

GS fluctuation has been shown to correlate with run-level summary metrics of head motion, cardiac and respiratory variables across subjects (Power et al., 2017). To explore if these factors might mediate observed time of day effects on GS fluctuation, we correlated twelve summary metrics (across domains of head motion, cardiac and respiratory motion) to GS fluctuation (Figure 2A) and to time of day (Figure 2B) across subjects for each run.

**Figure 2.**
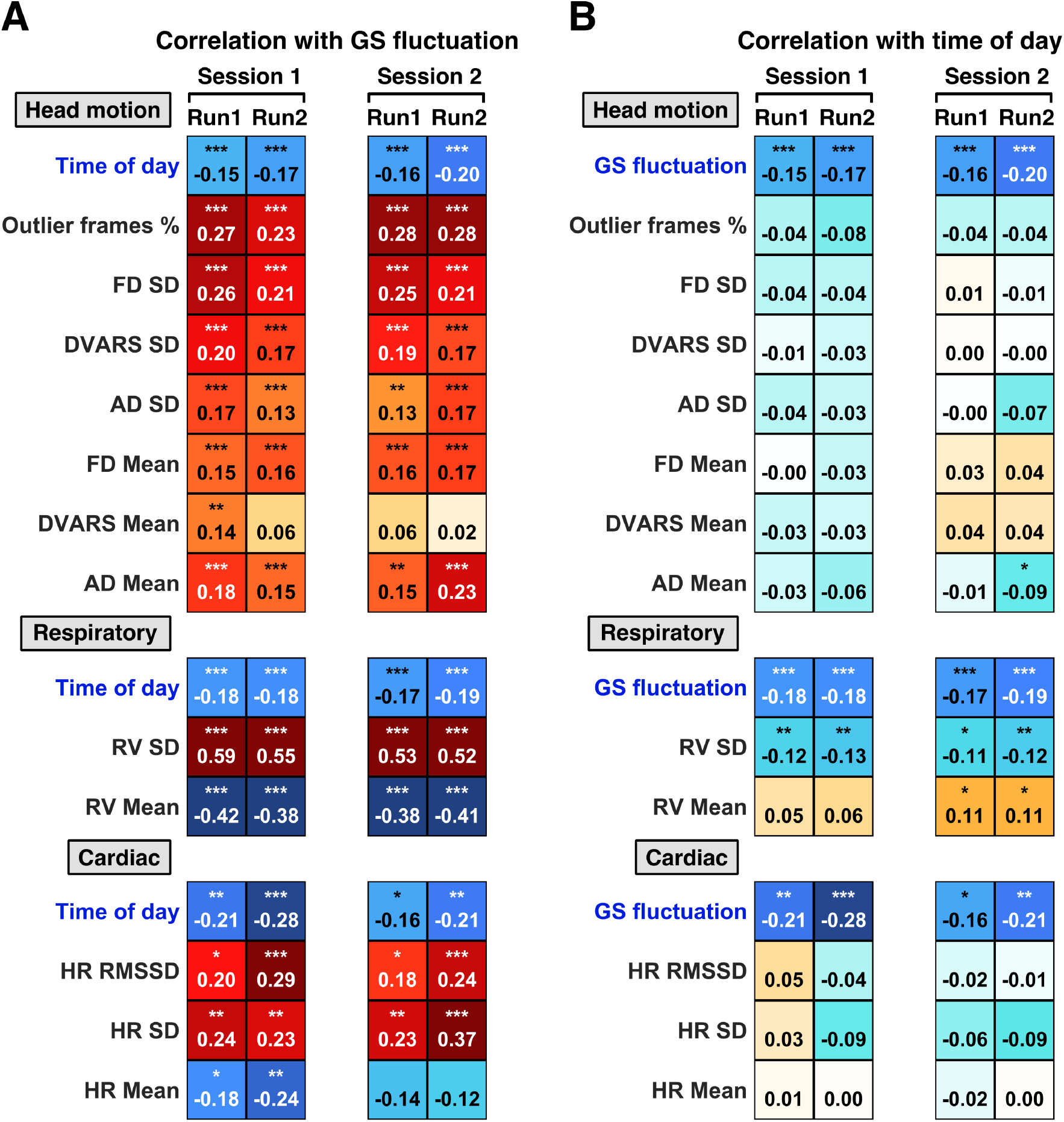
Head motion, respiratory and cardiac measures are strongly correlated with global signal (GS) fluctuation, yet only respiratory variation shows association with time of day. **(A)** Between-subject correlations of twelve run-level summary metrics with GS fluctuation. **(B)** Between-subject correlations of twelve run-level summary metrics with time of day. Due to exclusion of subjects with poor physiological data quality, different subgroups of subjects were used for analyses of head motion (Session 1: N = 942, Session 2: N = 869), respiratory (Session 1: N = 741, Session 2: N = 668) and cardiac measures (Session 1: N = 273, Session 2: N = 272). Correlation between GS fluctuation and time of day was repeated in each subgroup. Numbers denote z-scored Pearson r correlation coefficients. Stars indicate significant correlations following FDR correction (*q < 0.05; ** q < 0.01; *** q < 0.001). SD: standard deviation; AD: absolute displacement; FD: framewise displacement; RV: respiratory variation.

Some participants had to be excluded because of poor cardiac or respiratory recording quality (see Methods). Therefore, we ended up with different subgroups of participants for the head motion, cardiac and respiratory analyses. Consequently, we replicated the negative correlations between GS fluctuation and time of day for each subgroup (Figure 2A-B).

After correction for multiple comparisons (see Methods), eleven of the twelve measures showed consistent correlations with GS fluctuation across all four runs, with the exception of DVARS Mean, which was significantly correlated with GS fluctuation in only one of the runs (Figure 2A). The strongest correlations with GS fluctuation were observed for standard deviation of respiratory variation (RV SD; average r = 0.55 across four runs), mean of respiratory variation (RV Mean; average r = - 0.40), standard deviation of heart rate (HR Mean; average r = 0.27), outlier frames % (average r = 0.26) and standard deviation of framewise displacement (FD SD; average r = 0.23).

However, when we correlated these same twelve measures to time of day (Figure 2B), only RV SD showed a significant correlation with time of day for all four runs (average r = −0.12) after correction for multiple comparisons (see Methods). It is worth noting that the strength of this correlation was weaker than the correlation between GS fluctuation and time of day in the same subgroup of subjects (average r = −0.18).

### 3.4 Effects of time of day on GS fluctuation persist after correcting for respiratory variation

Given that both respiratory variation standard deviation (RV SD) and GS fluctuation were robustly correlated with time of day (Figure 2B), we tested if the effect of time of day on GS fluctuation remained significant after controlling for RV SD.

We found that RV SD was negatively correlated with time of day both between-subjects and within-subjects in both sessions in the subgroup of subjects with high quality respiratory data (Figures 3A-B and Figure S2). Like GS fluctuation, RV SD was elevated in the second versus the first run of each session (Figure 3C and Figure S2).

**Figure 3.**
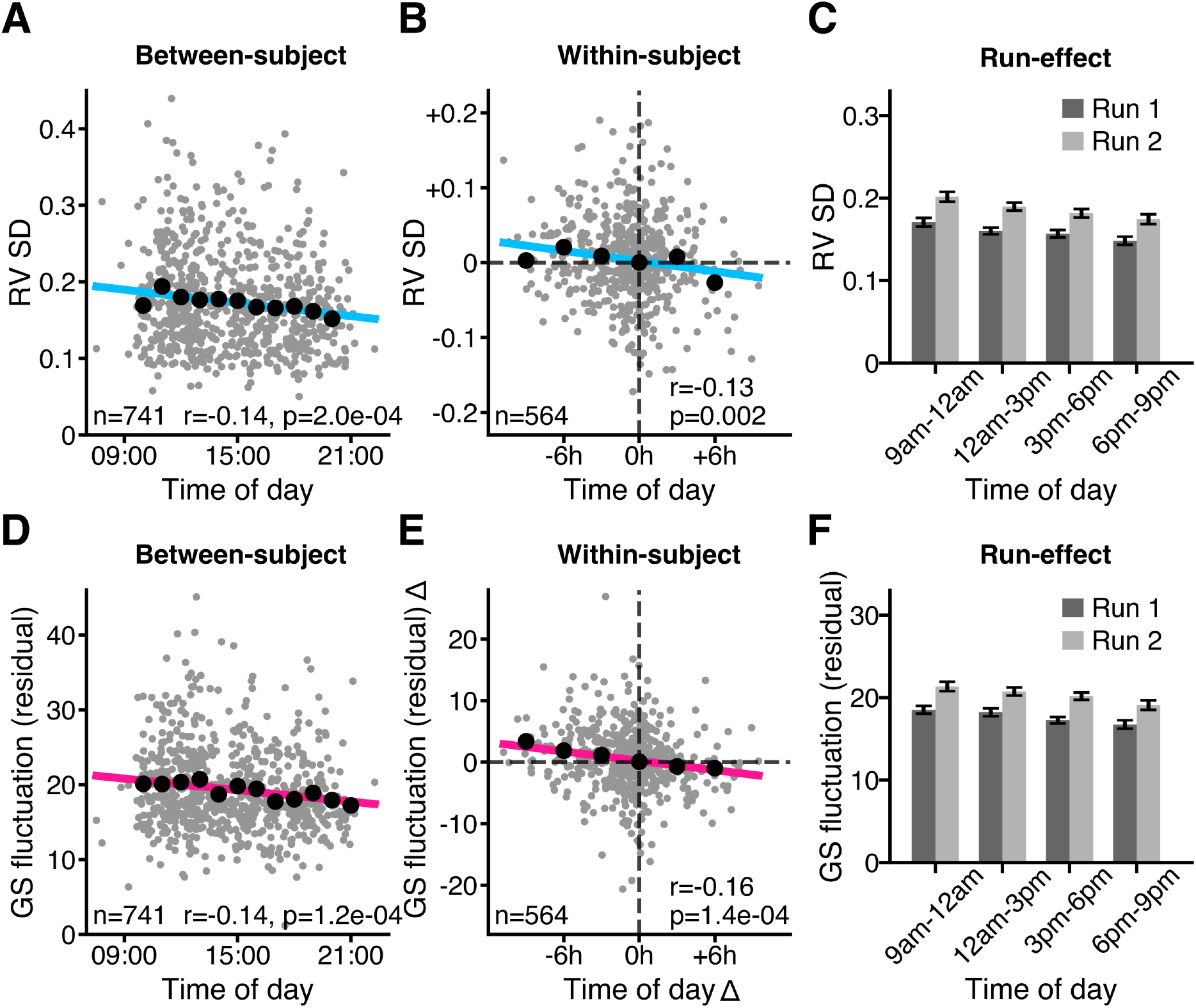
Negative association between time of day and GS fluctuation remains significant after controlling for respiratory variation. **(A)** Between-subject variation, (B) within-subject variation, and **(C)** run-effects of respiratory variation standard deviation (RV SD) as a function of time of day. **(D)** Between-subject variation, **(E)** within-subject variation and **(F)** run-effects of GS fluctuation residual (after regressing RV SD) as a function of time of day. Same as in Figure 1, within-subject effects were computed by taking the difference (Δ) for each variable between Session 2 and Session 1. Grey dots denote individual subjects. Black dots denote mean of GS fluctuation in hourly (**A, D**), or 3-hourly (**B, E**) time windows. r values denote Pearson r correlation coefficients. P values were derived from 100,000 permutations while keeping family structure intact (Winkler et al., 2015). SD refers to standard deviation. This figure shows the results for Session 1 (see Figure S2 for Session 2).

We then repeated our original analyses (Figure 1) after regressing RV SD from GS fluctuation across subjects. The negative correlation between time of day and GS fluctuation residual remained significant, and only slightly attenuated in magnitude (Figure 3D-F and Figure S2).

### 3.5 Time of day effects on regional BOLD signal fluctuation are most prominent in sensory-motor regions

The significant negative correlation between time of day and global signal fluctuation suggests that a substantial portion of the cortex might exhibit diurnal variation in activity during resting state. However, since the contribution of different regions to the global signal is often found to be non-uniformly distributed across the brain (Fox et al., 2009; Gotts et al., 2013; Power et al., 2017), we wanted to directly investigate the effects of time of day on brain-wide regional BOLD signal fluctuation.

We considered a whole brain parcellation consisting of 400 cortical regions (Figure 4A; Schaefer et al., 2018) and 19 subcortical regions (Figure 4B; Fischl, 2012). Regional BOLD signal fluctuation maps were derived for each subject by computing the standard deviation of the mean time series in each of the 419 regions of interest (ROIs) within each run. These run-level estimates were averaged within each session. We then correlated the regional BOLD signal fluctuation maps with time of day between-subjects for each session.

**Figure 4.**
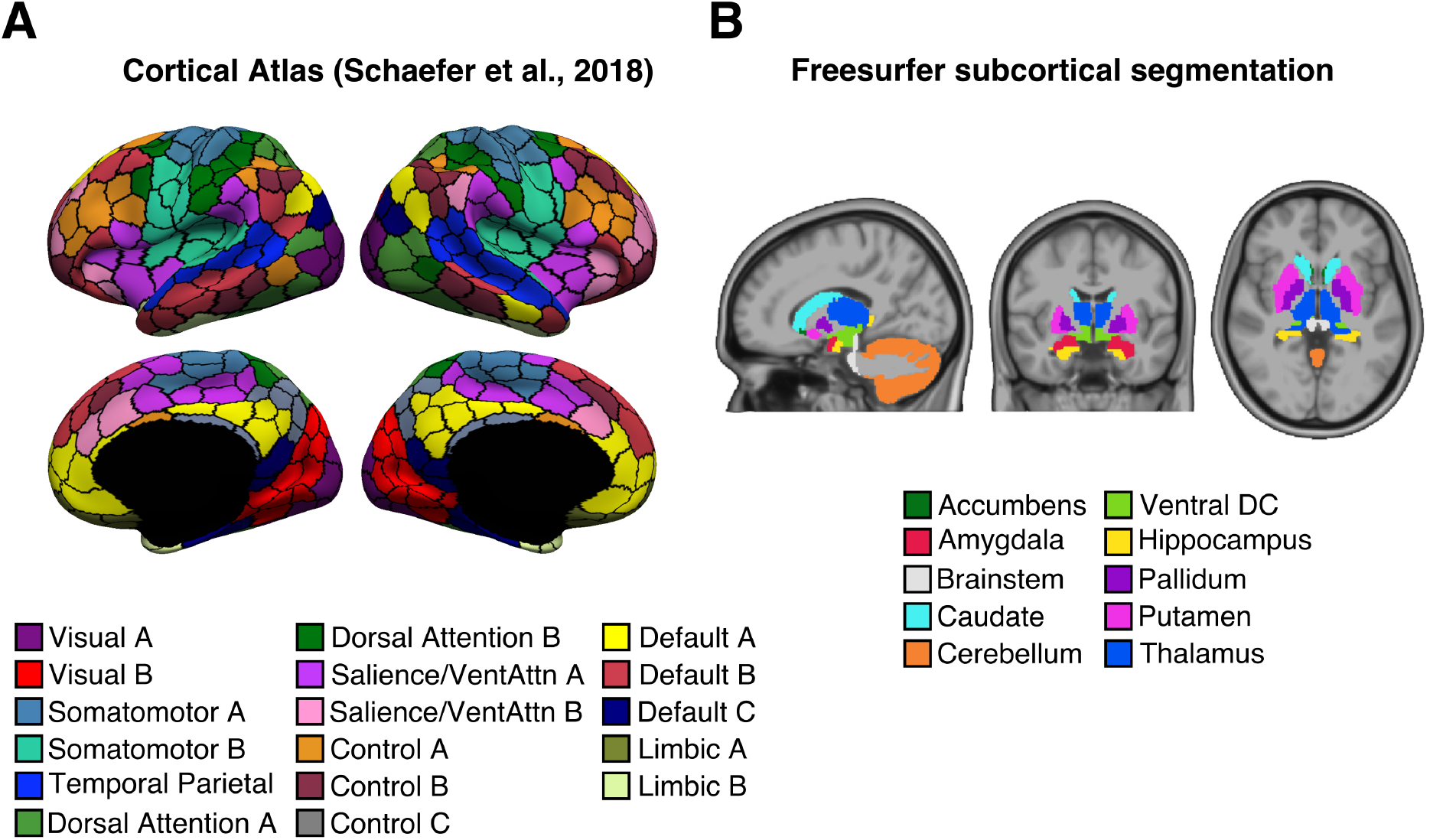
419 regions of interest (ROIs) **(A)** 400-area cortical parcellation in fs_LR surface space (Schaefer et al., 2018). Parcel colours correspond to 17 large-scale networks (Yeo et al., 2011). **(B)** 19 subcortical regions defined in subject-level volume space (Fischl et al., 2002).

Regional BOLD signal fluctuation showed a negative association with time of day in sensory cortices, including visual, somatomotor cortex, and in the anterior cingulate for Session 1 (Figure 5A). The impact of time of day was generally stronger and more widely distributed for Session 2, though again strong effects were observed in visual and somatomotor regions.

**Figure 5.**
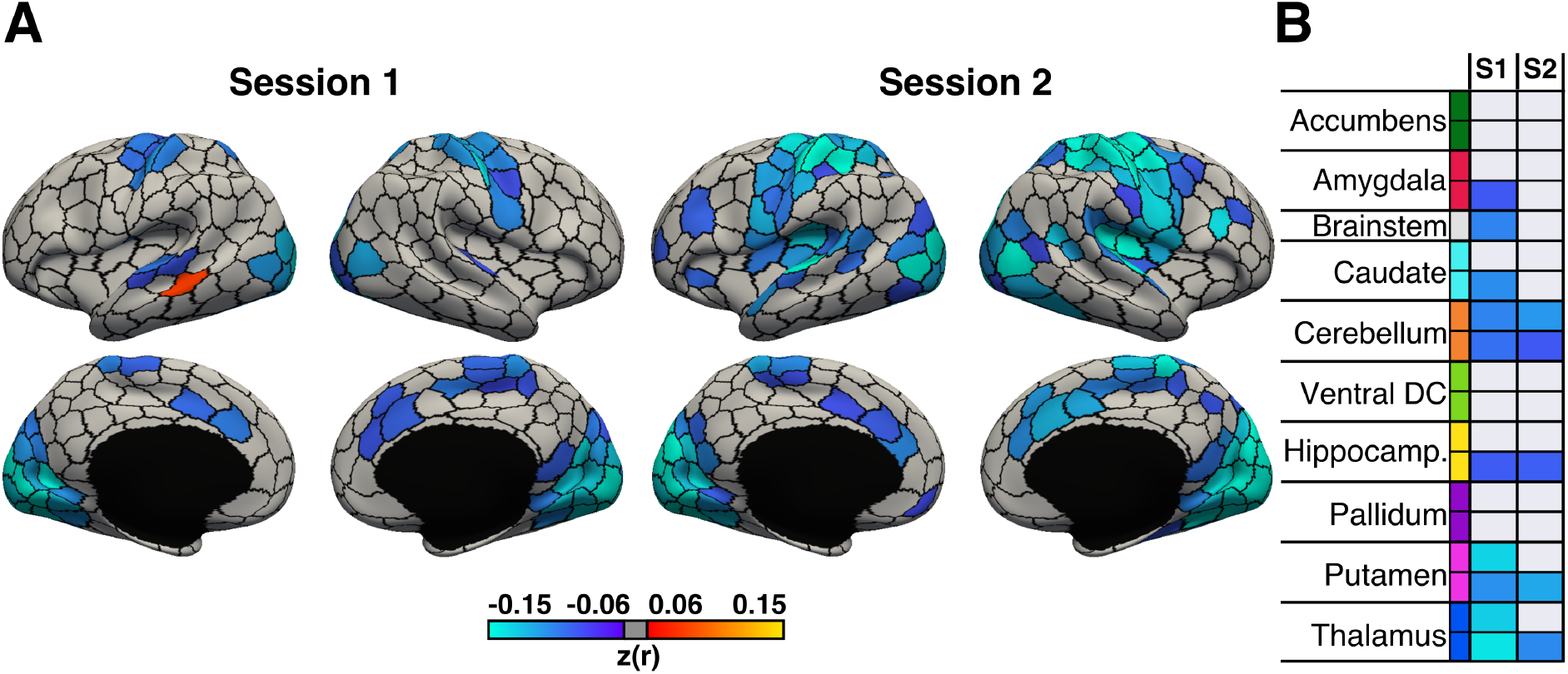
BOLD signal fluctuation is negatively correlated with time of day across cortical and subcortical regions during Session 1 (n = 942) and Session 2 (n = 869). Between-subject correlations between time of day and BOLD signal fluctuation for **(A)** 400 cortical and **(B)** 19 subcortical regions of interest. With the exception of brainstem, all subcortical regions are bilateral and presented as left-to-right hemisphere pairs (top to bottom). S1 denotes Session 1. S2 denotes Session 2. Colours (cool-warm) denote brain regions with significant Pearson r coefficients (q < 0.05, FDR-corrected), while non-significant regions are shown in grey. P values were derived from 100,000 permutations while keeping family structure intact (Winkler et al., 2015).

Similarly, subcortical regions exhibited either no significant association or a significant negative association between time of day and regional BOLD signal fluctuation (Figure 5B). The strongest effects across the subcortex were found in the bilateral putamen and bilateral thalamus during Session 1, effects that were replicated in the right hemisphere during Session 2 (Figure 5B).

### 3.6 Time of day is associated with widespread reductions in RSFC

There is significant interest in relating RSFC to individual differences in stable traits (Hampson et al., 2006; Van Den Heuvel et al., 2009; Finn et al., 2015; Li et al., 2019), an endeavour that is hindered by unaccounted variation in the behavioural or physiological states of subjects. This motivated us to assess whether variation in time of day of scans had a systematic effect on estimates of RSFC across subjects.

Like before, we computed a 419 x 419 correlation matrix for each subject, reflecting RSFC between 419 ROIs spanning the cortex and subcortex (Figure 4), using the ICA-FIX denoised data from each run. Then, session-level RSFC matrices were derived by averaging the two run-level RSFC matrices within each session. Finally, we computed the correlation between time of day and the RSFC of each ROI pair across subjects for each session. Results were visualised both at the region level (lower triangular) and at the network level (upper triangular). Network level results were computed by averaging Pearson r values across ROI pairs assigned to the same 17 large-scale networks defined in previous work (Yeo et al., 2011; Schaefer et al., 2018).

Time of day exhibited a negative correlation with RSFC throughout the brain (Figure 6A). Although the magnitude of time of day effects varied across networks and regions, this variation was highly consistent between the two sessions (See Figure S3 for Session 2). Prominent time of day effects were observed within Somatomotor and Visual networks, and between these two and Dorsal Attention and Default C network regions. In contrast, within-network RSFC in Default B, Control A & B and Limbic had minimal or no association with time of day.

**Figure 6.**
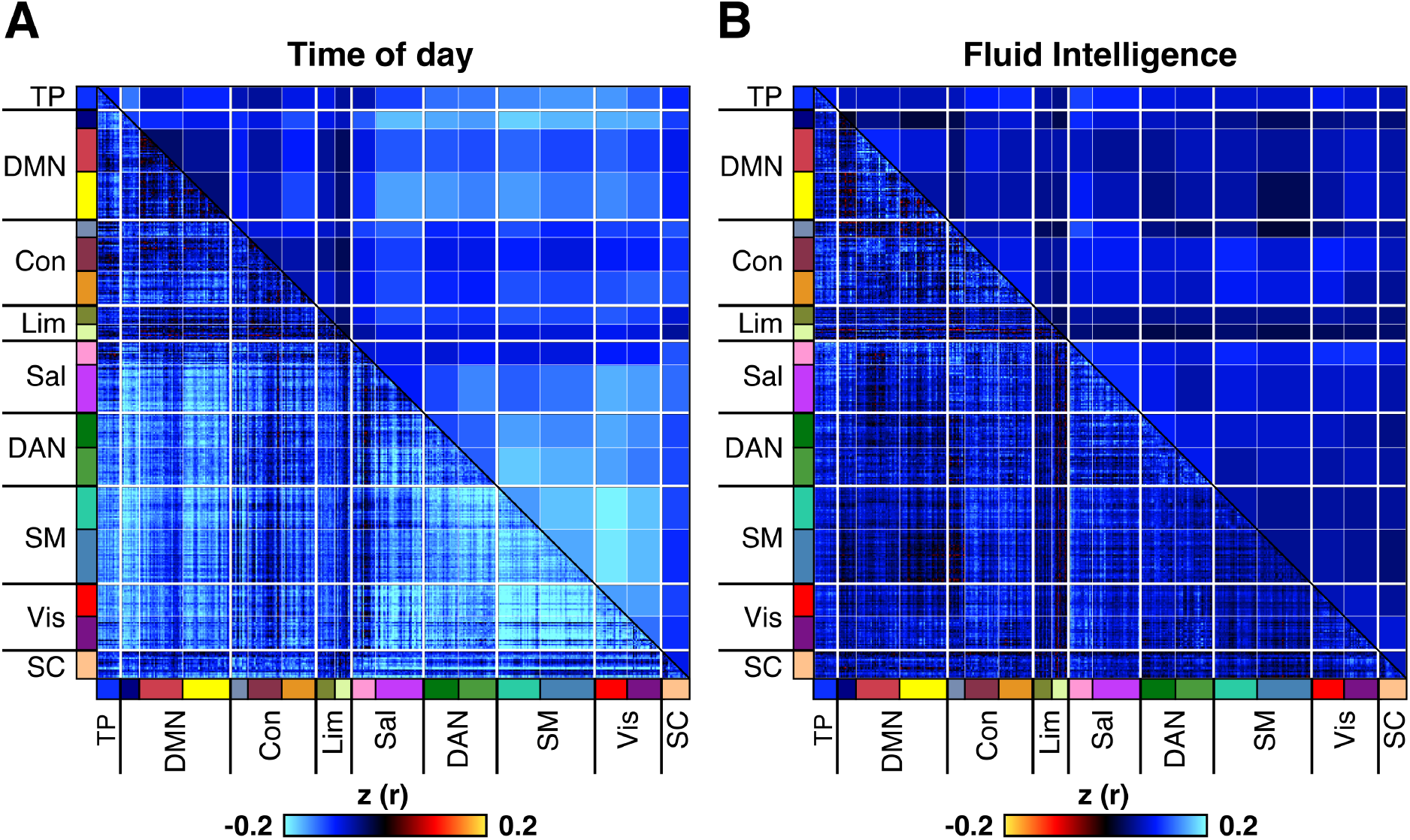
Resting state functional connectivity (RSFC) is negatively correlated with time of day across subjects, with a magnitude surpassing the strength of correlation between fluid intelligence and RSFC. Results shown are for Session 1 (n = 937; see Figure S3 for Session 2). **(A)** Correlation between time of day and RSFC across subjects. **(B)** Correlation between fluid intelligence and RSFC across subjects. Colours in lower triangular of correlation matrix denote Pearson r correlation coefficients. Colours in the upper triangular denote r values from the lower triangular averaged within network pairs. Colours on label axes denote correspondence of 419 regions to 17 large-scale cortical networks and to subcortex (SC). Association between time of day and RSFC was significant in both sessions as assessed by network-based statistics (FDR-corrected at q < 0.05), while association between fluid intelligence and RSFC association was only significant in Session 1 (FDR-corrected at q < 0.05). See Methods for details of network-based statistics (Zalesky et al., 2010). Note that the colour scale for fluid intelligence - RSFC correlation map was inverted to facilitate visual comparison with time of day effects. Fluid intelligence was chosen because it is a widely studied measure amongst resting-fMRI studies of brain-behavioural associations (Finn et al., 2015) and is one of the behavioural measures that is best predicted by resting-fMRI (He et al., 2018; Kong et al., 2019; Li et al., 2019).

To illustrate the magnitude of these effects, we also computed the correlation between RSFC and fluid intelligence. Fluid intelligence was chosen because it is a widely studied measure amongst resting-fMRI studies of brain-behavioural associations (Finn et al., 2015) and is one of the behavioural measures best predicted by resting state fMRI (He et al., 2018; Kong et al., 2019; Li et al., 2019). The correlation between time of day and RSFC was visibly and quantitatively stronger (Figure 6A; median absolute z = 0.13) than the correlation between fluid intelligence and RSFC (Figure 6B; median absolute z = 0.07) in Session 1. These findings were replicated in Session 2 (Figure S3).

### 3.7 Global signal regression weakens time of day effects, but introduces novel effects in specific networks

Global signal regression (GSR) is a widely used denoising method, effective at removing widely distributed (e.g. respiratory) artefacts from resting state fMRI data (Satterthwaite et al., 2013; Power et al., 2014; Burgess et al., 2016; Ciric et al., 2017; Power et al., 2017, 2018; Parkes et al., 2018; Li et al., 2019). However, GSR remains highly controversial because GSR might discard neural signals (potentially related to arousal) and introduce biases into the data (Murphy et al., 2009; Saad et al., 2012; Murphy and Fox, 2017; Glasser et al., 2018).

Since we identified a robust association between time of day and global signal fluctuation, we hypothesized that GSR might diminish time of day effects on regional BOLD signal fluctuation and RSFC. Thus, we repeated our original analyses of regional BOLD signal fluctuation and RSFC, but this time using time-series that had undergone GSR (see Methods).

GSR reduced the number of regions showing significant negative association between time of day and BOLD signal fluctuations (Figure S4). However, GSR also introduced positive correlations in several regions that previously showed no significant association with time of day (Figure S4).

GSR also resulted in an overall reduction of time of day effects on RSFC (Figure S5; median absolute z values in Session 1: z = 0.04, Session 2: z = 0.04) relative to our original analyses without GSR (Figures 6A and S3; median absolute z values in Session 1: z = 0.12, Session 2: z = 0.13). Nevertheless, the effect of time of day on RSFC remained significant in both sessions following GSR as assessed by network-based statistics (FDR-corrected at q < 0.05). However, GSR also amplified the association between time of day and RSFC in certain large-scale circuits, e.g., Somatomotor A – Control and Somatmotor A – Salience B (Figure S5).

## 4. Discussion

### 4.1 Summary

Despite widespread evidence from animal (Vyazovskiy et al., 2008; Gilestro et al., 2009; Liu et al., 2010) and human studies (Schmidt et al., 2009; Blautzik et al., 2013; Shannon et al., 2013; Huber et al., 2013; Muto et al., 2016; Cordani et al., 2018) that brain activity exhibits diurnal variation, most fMRI studies rarely consider the potential effects of time of day. In this study, we examined the impact of time of day on various measures of resting state brain activity in over 900 participants from the HCP dataset, scanned between 8 am to 10 pm on two sessions. We observed time of day dependent reductions in the magnitude of GS fluctuation, regional BOLD signal fluctuation, as well as reductions in whole-brain RSFC throughout the course of the day. We also demonstrated that the magnitude of whole-brain RSFC decrease was larger than the association between RSFC and fluid intelligence.

### 4.2 Time of day and arousal have opposing effects on GS fluctuation at distinct timescales

Surprisingly, the directionality of our main findings was in direct contrast to our hypothesis, which was motivated by models of circadian fluctuations in arousal levels (Strogatz et al., 1987; Van Dongen and Dinges, 2000; Bes et al., 2009; de Zeeuw et al., 2018). We hypothesized that the magnitude of GS fluctuation, a measure associated with drowsy states (Wong et al., 2013; Yeo et al., 2015; Liu et al., 2017) would be lowest in the late morning and early evening hours, when levels of arousal are typically at peak levels, and highest in the mid-afternoon when levels of arousal often dip. Instead, we observed the highest levels of GS fluctuation in the late morning hours, followed by a gradual reduction of GS fluctuation across subjects throughout the day until the late evening hours. Although the HCP resting state scan protocol did not involve concurrent assessment of arousal (e.g. Ong et al., 2015; Chang et al., 2016; Wang et al., 2016), it is unlikely that arousal levels would be lowest in the late morning and monotonically rise throughout the entire day.

However, we did observe a robust increase in GS fluctuation in the second run of each session. Similar within-session variation in the magnitude of BOLD signal fluctuation have been previously reported in the HCP (Bijsterbosch et al., 2017) and in other datasets (Tagliazucchi and Laufs, 2014). Furthermore, simultaneous resting state fMRI and polysomnographic EEG recordings have shown that subjects exhibit an increased propensity to fall asleep with longer duration of time spent in the scanner (Tagliazucchi and Laufs, 2014). These studies, as well as a wealth of earlier work linking higher regional BOLD signal (Kiviniemi et al., 2005; Horovitz et al., 2008; Bianciardi et al., 2009; Larson-Prior et al., 2009; Licata et al., 2013) and GS (Wong et al., 2013; Yeo et al., 2015; Liu et al., 2017) fluctuations to low arousal states, suggest that this phenomenon most likely reflects a drop in arousal as subjects become progressively more drowsy while lying in the scanner. Interestingly, we found that this between-run increase in GS fluctuation levels was present throughout different times of day in both sessions. Thus, we observed a run-level increase in GS fluctuation on the scale of minutes that was superimposed on a slower downward trend in GS fluctuation on the scale of hours.

### 4.3 Potential roles of homeostatic mechanisms and cerebral blood flow changes

What mechanism could account for the observed time of day associated reductions in GS fluctuation and RSFC, if not circadian variation in arousal levels as we had hypothesised originally? A growing number of animal studies have suggested that wakefulness and sleep might be respectively associated with homeostatic upscaling and downscaling of synaptic strength (Tononi and Cirelli, 2003; Vyazovskiy et al., 2008; Gilestro et al., 2009; Liu et al., 2010). Building on this work, human studies have found that cumulative wakefulness is associated with a gradual rise in cortical excitability, as measured by TMS-evoked EEG potentials (Huber et al., 2013; Kuhn et al., 2016) and motor-evoked potentials (Kuhn et al., 2016). Interestingly, in one of these studies, the increase in cortical excitability was detectible even prior to sleep deprivation and was not associated with concomitant changes in behavioural indices of arousal (Kuhn et al., 2016). However, it’s not directly apparent how a diurnal increase in synaptic strength or cortical excitability would result in the time of day-associated reductions in GS fluctuation and RSFC that we observed in our study.

Based on the premise that synaptic strengthening has a significant metabolic cost (Harris et al., 2012), the synaptic homeostasis hypothesis (Tononi and Cirelli, 2003) predicts an increase in cerebral oxygen and glucose consumption, and as a result, increased cerebral blood flow (CBF) with cumulative wakefulness (Tononi and Cirelli, 2006). Indeed, several studies have reported elevated CBF in the evening relative to morning (Braun, 1997; Kuboyama et al., 1997; Elvsåshagen et al., 2019; but see Shannon et al., 2013; Hodkinson et al., 2014).

Interestingly, the largest of these studies (Elvsåshagen et al., 2019) reported regional CBF increases related to duration of wakefulness in the same regions where we observed the strongest time of day-associated reductions in resting state BOLD signal fluctuation and RSFC (e.g. somatomotor and visual cortex). These same regions have also been shown to exhibit reduced task-related BOLD signal activation as a function of duration of wakefulness (Muto et al., 2016). This is of particular relevance, since elevation of baseline CBF levels via experimentally induced hypercapnia (inhalation of CO_2_ enriched air) is known to reduce the dynamic range of resting state BOLD fluctuation and RSFC (Biswal et al., 1997; Xu et al., 2011; Liu, 2013). Thus, a time of day associated increase in baseline CBF would present a possible mechanism that could account for our observed reductions in GS fluctuation and RSFC. Nevertheless, the jury is still out on whether these effects are indeed the consequence of homeostatic upscaling of synaptic strength or another mechanism.

### 4.4 Respiratory variation also shows time of day modulation but does not account for changes in GS fluctuation

Since respiration is a potent modulator of BOLD signal fluctuation (Wise et al., 2004; Birn et al., 2006; Power et al., 2018), we considered whether diurnal changes in respiratory variation existed, and whether these could account for the time of day associated reductions in GS fluctuation. Indeed, we observed a significant negative correlation between the standard deviation of respiratory variation (RV SD) and time of day. Thus, subjects scanned in the earlier hours of the day exhibited more dynamic breathing patterns (potentially more sighs, yawns, pauses, or variation in breathing rate or depth) than those scanned in the evening. The mechanisms behind this are unclear, although earlier studies have proposed that breathing stability might be impaired in the morning hours, as suggested by augmented ventilatory response to hypoxia (Gerst et al., 2011) and hypercapnia challenges (Cummings et al., 2007).

Variation in respiratory activity, whether spontaneous (Power et al., 2017, 2018) or instructed (Birn et al., 2006, 2008) is known to introduce variance into the BOLD signal, via CO_2_ associated fluctuations in CBF (Wise et al., 2004). Indeed, we observed a strong positive correlation between RV SD and GS fluctuation, replicating earlier findings from other datasets (Power et al., 2017, 2018). Furthermore, we found that the between-run elevation of GS fluctuation, was also accompanied by a between-run elevation of RV SD. Given the strong statistical relationship between the two measures, as well as a plausible mechanistic link (Wise et al., 2004), we thought that respiratory variation might account for the effect of time of day on GS fluctuation. However, the negative association between time of day and GS fluctuation remained significant and only slightly diminished following regression of RV SD from GS fluctuation. Thus, respiratory variation only accounted for a small component of the observed diurnal downward shift in GS fluctuation.

### 4.5 Spatial distribution of time of day effects

Time of day was negatively correlated with regional BOLD signal fluctuation and RSFC across many brain regions, with the strongest effects apparent in visual and somatomotor cortex. These findings are consistent with several observations (though not all) from previous smaller scale studies (Blautzik et al., 2013; Jiang et al., 2016; Cordani et al., 2018).

One of the first resting state fMRI studies to look at time of day effects found that within-network strength and spatial extent of the somatomotor network decreased as a function of time elapsed since participant’s midsleep hour, while executive control network connectivity was characterised as particularly stable over time (Blautzik et al., 2013). Later, Jiang and colleagues showed reduced amplitude of low frequency fluctuations and homogeneity in sensory-motor regions in evening versus morning scans (Jiang et al., 2016). More recently, Cordani and colleagues reported diurnal reductions in sensory-motor regions during morning and evening twilight hours, and that reductions of BOLD signal fluctuation in visual cortex were associated with improved performance on a visual detection task (Cordani et al., 2018). To our knowledge, ours is the first study to report a cumulative reduction of GS fluctuation, and a negative correlation between regional BOLD signal fluctuation and time of day.

A number of studies outside the context of investigating time of day effects, have shown that sensory-motor regions exhibit particularly high levels of intra-subject variation in BOLD signal fluctuation and RSFC across the brain (Mueller et al., 2013; Laumann et al., 2015; Bijsterbosch et al., 2017; Kong et al., 2019), an observation typically attributed to modulations of arousal state (Tagliazucchi and Laufs, 2014; Schneider et al., 2016; Bijsterbosch et al., 2017; Liu et al., 2018; Glasser et al., 2018).

However, Power and colleagues have cautioned that since regressors that account for respiratory signals also exhibit a strong loading on these same regions, the high levels of variability in these regions could reflect respiratory artefacts rather than neural activity related to arousal (Power et al., 2017; Power, 2019). Furthermore, they have drawn attention to the fact that there is a high prevalence of isolated deep breaths during resting state scans (Power et al., 2019). These large breaths produce BOLD signal modulations spanning 30-40 seconds that are far slower than the standard evoked response function used in task fMRI to model neural activity (Power et al., 2019).

The challenge of distinguishing neural fluctuations from potential respiratory artefacts is made difficult by the fact the respiratory and neural dynamics are unlikely to be independent even without the presence of artefacts. For example, a recent study found that natural fluctuations in end-tidal pCO_2_ due to respiration appeared to track oscillatory power in multiple frequency bands as measured by MEG (Driver et al., 2016).

Since, electrophysiological techniques, such as MEG and EEG, do not rely on neurovascular coupling, they provide a more direct measurement of neural activity without fMRI’s susceptibility to respiratory artefacts. Indeed, studies employing electrophysiology in tandem with fMRI have provided many cases in point where BOLD global signal dynamics vary together with neural activity as indexed by local field potentials (Scholvinck et al., 2010; Wong et al., 2013; Liu et al., 2015; Chang et al., 2016; Bergel et al., 2018).

Chang and colleagues reported that the global fMRI signal showed a strong temporal correlation with a local field potential (LFP) based index of arousal in macaques (Chang et al., 2016). In another study, Liu and colleagues found that global peaks observed in fMRI signal were associated with spectral shifts in LFP in macaques, and that peaks defined in the same manner exhibited a visual-sensorimotor co-activation pattern in humans (Liu et al., 2018). More recently, Bergel and colleagues used ultrasound imaging in rats to show that brain wide vascular surges lasting between 5 to 30 seconds during REM sleep were preceded by sustained activity in the theta, mid-gamma, and high-gamma LFP bands (Bergel et al., 2018). Importantly, Turchi and colleagues demonstrated that pharmacological inactivation of basal forebrain nuclei selectively decreased global fMRI signal fluctuation ipsilaterally to the injection site even after statistically accounting for end-tidal pCO_2_ fluctuation and subject motion (Turchi et al., 2018).

### 4.6 Head motion, heart rate variability and cerebrovascular reactivity

Recent work has shown that substantial individual variation in global signal fluctuation (Power et al., 2017, 2018) and RSFC (Van Dijk et al., 2012; Satterthwaite et al., 2012; Power et al., 2012; Geerligs et al., 2017) can be explained by summary metrics derived from recordings of physiological variables, even after thorough preprocessing of the data. Our findings corroborate this view to the extent that we also observed strong correlations between GS fluctuation and a wide range of summary metrics across domains of respiration, head motion and cardiac measures. Yet, when we correlated these same metrics with time of day, only RV SD was consistently correlated with time of day across the four runs in two sessions. Thus, individual differences in head motion and cardiac measures, while contributing to variation in GS fluctuation, did not account for the effects of time of day on resting brain activity.

Can time of day related variation in cerebrovascular reactivity explain our findings, whereby the same level of respiratory or neural activity might produce a stronger modulation of the BOLD signal at different times of day? Indeed, cerebrovascular reactivity has been shown to be reduced in the morning relative to evening hours, as indicated by lower by middle cerebral artery velocity and cerebral oxygenation following breath holds or hypercapnia challenges (Ameriso et al., 1994; Cummings et al., 2007; Ainslie et al., 2007). However, this account is not consistent with our observations of increased BOLD signal fluctuation and RSFC in the morning compared to evening, since lower cerebrovascular reactivity is typically associated with lower, not higher, BOLD signal fluctuation and RSFC (Golestani et al., 2016).

### 4.7 Implications for global signal regression

Our results suggest that while global signal regression (GSR) does not fully eliminate time of day effects on RSFC, it does lead to an overall reduction in time of day effects across the brain. Interestingly, the negative association between time of day and BOLD signal fluctuation remained strongly preserved in the visual cortex, in both sessions, even following GSR. This suggests that the visual cortex might be particularly susceptible to diurnal variation, an idea that is consistent with recent task-activation studies (Cordani et al., 2018; Muto et al., 2016).

The validity of GSR has been extensively debated in the field of resting state fMRI (Murphy and Fox, 2017). Proponents of GSR have emphasised its ability to remove spatially widespread respiratory artefacts (Power et al., 2014, 2017, 2018; Power, 2019), while others have raised concerns that this may entail removal of neural signal (Gotts et al., 2013; Yang et al., 2014; Glasser et al., 2018, 2019) or the introduction of spurious relationships into the data (Murphy et al., 2009; Saad et al., 2012; Satterthwaite et al., 2012). Our control analyses suggested that the effect of time of day on GS fluctuation could not be accounted for by individual differences in measures that might suggest non-neural sources of variance, e.g. head motion, respiratory and cardiac measures. Thus, assuming that our results indeed reflect diurnal variation in large-scale spontaneous brain activity, then the current study could represent an example of where GSR might be discarding signal of neural origin.

Whether, GSR is preferable or not, however still might vary according to the dataset at hand (e.g. extent of global artefacts in the data) and on the specific aims of the study. For example, it is possible that time of day effects on resting brain activity might simultaneously manifest as both globally distributed decreases in RSFC, and as more subtle, localized increases of RSFC between a few select regions or networks. These latter effects, however, might only become apparent once the global signal is removed from the data, for example, as was the case with the positive correlations between time of day and Somatomotor – Control network RSFC in the current study (Figure S5).

Indeed, there has been substantial interest in the field to pioneer alternatives to global signal regression (Behzadi et al., 2007; Chai et al., 2012; Power et al., 2018; Glasser et al., 2018), with the aim of removing putative global artefacts, without some of the purported drawbacks of GSR, such as the inadvertent removal of neural signals of interest. Nevertheless, the validity and utility of various approaches for dealing with global signals remains a topic of debate and enquiry (Murphy and Fox, 2017; Glasser et al., 2018; Power, 2019; Glasser et al., 2019; Li et al., 2019).

### 4.8 Methodological limitations

Since our main finding of a time of day associated reduction in GS fluctuation was derived based on group-level analysis, we cannot confidently infer what proportion of individuals actually exhibited this tendency within our sample. It is possible that for a subset of our sample there was no effect of time of day on GS fluctuation, or an effect with a different pattern than the one observed across the group.

The HCP protocol did not involve assessment of arousal state during the resting state scans. Methods typically used in the field include tracking of eyelid closures (Ong et al., 2015), pupillometry (Schneider et al., 2016), EEG (Wong et al., 2013), polysmonography (Tagliazucchi and Laufs, 2014) or cognitive vigilance (Muto et al., 2016). Such data would have helped interpret the unexpected directionality of our findings by providing a more direct measure of arousal state, e.g. changes in the frequency of spontaneous eye-closures or duration of in-scanner sleep. Similarly, measurement of spontaneous fluctuations in end-tidal pCO_2_ levels via a nasal cannula during the resting state scans (Wise et al., 2004) would have enabled a more comprehensive characterisation of the effects of time of day on respiration, and the extent to which these effects can potentially account for diurnal variation in GS fluctuation.

Our ability to detect an association between time of day and GS fluctuation across subjects had as much to do with a sufficiently low inter-subject variation in GS fluctuation, as with a sufficiently large effect of time of day on GS fluctuation. For example, our inability to detect time of day effects on head motion or heart rate variability measures might have been due to greater baseline inter-individual variation of these measures relative to the magnitude of the impact of time of day. Thus we cannot rule out the existence of time of day effects for these other measures.

Finally, there are several unaccounted variables which could have potentially impacted the observed diurnal variation in resting brain activity. In the current study, time of day likely correlates with the hours of scan usage on a given day, thus we cannot rule out the presence of scanner use-related hardware artefacts. However, we note that such hardware artefacts are very unlikely to produce time of day effects with a spatial specificity to visual and somoatomotor cortices. Although caffeine consumption was not monitored, increased morning caffeine consumption would not account for elevated GS fluctuation, since caffeine is known to reduce GS fluctuation (Wong et al., 2012). Other examples of unaccounted variables include meal timing, meal size and composition, fluid intake, bedtime on previous night, wakeup time on the day of study, timing of commute, weather and light levels.

### 4.9 Recommendations

We believe our findings build a strong case for the importance of accounting for time of day, based on having revealed systematic time of day effects on resting state fMRI measures greater in magnitude to behavioural associations (e.g. fluid intelligence), consistent in both between- and within-subject analyses, present following regression of respiratory variation, and replicable across two sessions within a widely used, open-access, large-scale dataset. In light of these findings we would like to make a number of recommendations.

1. We suggest reporting time of day of fMRI scans and of other experimental protocols and measurements. Even if all subjects were scanned at the same timeslot within a particular study, reporting time of day could help account for between-study variation in results and potentially even failed replications. Meta-analyses could then be leveraged to explore how time of day affects various regions, networks and tasks across different domains of the literature.
2. In individual studies where subjects are scanned at different times, we suggest assessing the impact of time of day on main effects of interest, and considering group-level correction for time of day. Large-scale studies, such as the HCP, are particularly vulnerable to time of day effects as subjects are often scanned throughout the day to speed up data collection.
3. Since time of day effects might reflect the impact of duration of wakefulness on brain states, we encourage assessment and reporting of sleep patterns of subjects in the preceding weeks and on the day of study, via self-report and via implicit methods such as actigraphy. This is already standard practice in studies investigating circadian mechanisms (Muto et al., 2016; Cordani et al., 2018). However it could be potentially beneficial to extend to all studies that examine individual variation in behavioural traits that could be affected by circadian phase, duration of wakefulness or fatigue.
4. Case-control studies that examine clinical populations with altered sleep duration or circadian phase (e.g. schizophrenia, bipolar disorder, elderly) could be affected by individual differences in duration of wakefulness or circadian phase, even if all scans take place at the same time of day. For these studies it might make sense to match control and patient groups for duration of wakefulness at the time of scanning.

## 5. Materials & Methods

### 5.1 Data Overview

#### 5.1.1 Subject characteristics

All data analysed in this manuscript is from the S1200 release of the Human Connectome Project (HCP) dataset (Van Essen et al., 2012; Smith et al., 2013), which was drawn from a population of young, healthy twins and siblings. The scanning protocol, subject recruitment procedures and informed written consent forms, including consent to share de-identified data, were approved by the Washington University institutional review board. Subjects underwent a range of neuroimaging protocols, including resting state fMRI, which was collected over two sessions, with each session split into two consecutive runs. We only considered subjects who had available resting state fMRI data from at least one of the two sessions, acquired with both left-to-right and right-to-left phase encoding. We excluded from analysis, any session where the subject exhibited excessive head motion in more than 50% of the acquired frames in either run. Following exclusion of excessive movers, we retained 942 subjects from Session 1 and 869 subjects from Session 2. We conducted separate, parallel analyses for each session, with the exception of the within-subject correlations, which required analyses of subjects with data from both sessions (N = 831). Additional exclusions were applied selectively for analyses of cardiac and respiratory data based on the quality of recordings and for analyses of RSFC due to a handful of subjects missing fluid intelligence scores (see below).

#### 5.1.2 Image acquisition

Imaging took place at Washington University on a custom-made Siemens 3 T Skyra scanner. Four runs of resting state fMRI data were collected over two sessions (2 on each session), which for the majority took place on separate days. During each run 1200 frames were acquired using a multi-band sequence at 2 mm isotropic resolution with a TR of 0.72 seconds over the span of 14.4 minutes. Subjects were instructed to maintain fixation on a bright cross-hair presented on a dark background in a darkened scanning room. The resting state sequences were collected at the start of each scanning session, preceded only by brief field-map and BIAS acquisitions. Structural images were acquired at 0.7 mm isotropic resolution. Further details of the data acquisition parameters can be found in previous publications (Van Essen et al., 2012; Smith et al., 2013).

#### 5.1.3 Preprocessing of structural images

Preprocessing of T1 structural images was implemented using FreeSurfer (Fischl, 2012). This included brain extraction, subject-level volumetric segmentation of subcortical regions, cortical surface reconstruction and registration of each subject’s cortical surface mesh to a common spherical coordinate space (Fischl et al., 1999a, 1999b).

#### 5.1.4 Preprocessing of resting state fMRI

The HCP release includes resting-state fMRI data at two stages of preprocessing: minimally preprocessed and spatial ICA-FIX cleaned data (Glasser et al., 2013). The minimally preprocessed data had undergone removal of spatial distortion, motion correction via volume re-alignment, registration to the structural image, bias-field correction, 4D image intensity normalization by a global mean, brain masking, volume to surface projection, multimodal inter-subject alignment of the cortical surface data (Robinson et al., 2014), and 2 mm (FWHM) parcel- and surface-constrained smoothing. The resulting image is a CIFTI time-series with a standard set of grayordinates at 2 mm average surface vertex and subcortical volume voxel spacing. The HCP release of the spatial ICA-FIX processed data (ICA-FIX) had undergone additional preprocessing steps following minimal preprocessing. The data was first de-trended with a temporal high-pass filter and corrected for head-motion via a 24 parameter regression, before undergoing denoising via spatial ICA-FIX (Griffanti et al., 2014; Salimi-Khorshidi et al., 2014).

#### 5.1.5 Statistical analyses

Given the high prevalence of families in the dataset, we applied a method of permutation testing, which respects the family-structure of participants during shuffling (Winkler et al., 2015). P-values were calculated from 100,000 permutations.

### 5.2 Time of day effects on global signal fluctuation

We assessed the relationship between time of day and global signal fluctuation between-subjects in each session, followed by a within-subject (i.e. between-session) assessment of the same relationship. The between-subject association between time of day and global signal fluctuation was computed for each session, by first averaging time of day and global signal values across the two runs for each subject, followed by computing a Pearson r correlation coefficient between the two session-level average values. For the computation of within-subject correlation, we again averaged each measure across the two runs of each session for each subject, and then used the within-subject difference (Δ) of each measure between the two sessions to compute a Pearson r correlation.

### 5.3 Assessment of non-neural contributions to time of day effects

#### 5.3.1 Head motion, cardiac and respiratory summary metrics

We also examined the impact of time of day on a range of head motion, respiratory and cardiac variables, which have been highlighted in the literature as potential indicators of non-neural sources of variance. In each of the four runs, we calculated 12 subject-level summary metrics: percentage of frames that are outliers (Outlier frames %), standard deviation of Framewise Displacement (FD SD), mean of Framewise Displacement (FD Mean), standard deviation of Absolute Displacement (AD SD), mean of absolute displacement (AD Mean), standard deviation of DVARS (DVARS SD), mean of DVARS (DVARS Mean), standard deviation of Respiratory Variation (RV SD), mean of Respiratory Variation (RV Mean), root-mean of the successive differences of heart rate (HR RMSSD), standard deviation of instantaneous heart rate (HR SD), mean of heart rate (Mean HR).

Outlier frames were defined as those where DVARS exceeded 75 or where FD exceeded 0.2 mm, as well as frames one before and two after such outlier frames, and non-outlier segments consisting of fewer than five consecutive frames. This definition is based on earlier work on head-motion censoring (Power et al., 2012, 2014), while the specific parameters are consistent with our recent study using the HCP dataset (Kong et al., 2019). FD was defined as the summed root-mean-square of the derivatives of six motion parameters (3 rotational, 3 translational), AD was defined as the summed root-mean-square of six motion parameters (3 rotational, 3 translational), and DVARS was defined as the root-mean-square of voxelwise differentiated signal. FD, AD and DVARS were all computed using the FSL function *fsl_motion_outliers* (Jenkinson et al., 2012). Respiratory variation (RV) was defined as the standard deviation of the respiratory waveform within 5.76 second windows (equal to 8 fMRI frames) centred on each fMRI frame (Chang et al., 2009). HR was calculated at each peak based on the preceding inter-beat-interval.

We correlated (Pearson r) each of the 12 summary metrics separately with both global signal fluctuation and with time of day across subjects. Our intent for correlating them with global signal fluctuation was to provide us with a positive control, as many of these measures have been shown to be associated with global signal fluctuation in other datasets (Power et al., 2017). Significance level was calculated using 100,000 permutations, while respecting family structure of participants, and FDR-corrected at q < 0.05.

#### 5.3.2 Cohorts for analysis of respiratory and cardiac measures

All pulse and respiratory time-series in the S1200 HCP dataset underwent visual inspection by the same person (CO). For a subject’s data to be included in the analysis of a session, both runs had to pass quality control. Following quality control, a substantial proportion of the pulse data, and a somewhat lesser proportion of the respiratory data, were considered to be inadequate for analysis. Therefore, separate subgroups were used for analysis of head motion (Session 1: N = 942; Session 2: N = 869), respiratory (Session 1: N = 741; Session 2: N = 668) and cardiac summary metrics (Session 1: N = 273; Session 2: N = 272).

#### 5.3.3 Quality control of pulse data

We used the pulse oximeter and respiratory waveform data that was acquired (at 400 Hz) during the resting state fMRI. We applied automated peak detection on the pulse wave data (https://github.com/demotu/BMC/blob/master/functions/detect_peaks.py), followed by individual visual inspection of each peak, and manual correction where necessary. Runs containing peaks that could not be reliably identified due to possible motion or hardware artefacts were excluded. Isolated ectopic beats were manually deleted, but were not a reason for excluding an entire run. In rare cases, runs with clearly identifiable peaks were excluded if they contained extended segments that were deemed to be arrhythmic or artefactual in origin (Figure S7). Our aim was not to classify arrhythmias in the HCP dataset, but to exclude runs where abnormal heart rhythms might inflate summary measures of heart rate variability, which under healthy sinus rhythm reflect the influence of the autonomic nervous system (Quintana et al., 2016). In a number of runs, subjects exhibited a heart rate of exactly 48.0 beats per minute with close to 0 inter-beat interval variability. These runs could be easily identified as outliers on Poincaré plots or on heart rate versus heart rate variability scatter plots (Figure S7), and thus were excluded them from further analysis.

#### 5.3.4 Quality control of respiratory data

The raw respiratory waveform data of each subject from each run was z-scored, lowpass filtered at 2 Hz, visualized in its entirety and then each run was manually checked for sufficient data quality. RV, our primary measure of interest, was defined as the standard deviation of the filtered respiratory waveform within 5.76 second sliding windows (equivalent to 8 TR), and plotted alongside the respiratory waveform data. The respiratory waveform frequently contained trains of high-frequency artefacts. However, in most cases these artefacts did not lead to changes in RV, as RV indexes fluctuations on a slower timescale that is calibrated to measure breathing patterns. We found the lowpass filter to be effective at eliminating many of these high frequency artefacts, without distorting RV in clean segments of data. Runs with poor signal quality, ambiguous waveforms, or sustained high frequency artefacts that were reflected in RV even after bandpass filtering, were excluded from analysis.

### 5.4 Time of day effects on regional BOLD signal fluctuation

To assess the spatial distribution of time of day effects across the brain, we computed, the Pearson’s r correlation between the time of day of each subject’s scan and the corresponding regional BOLD signal fluctuation measured across 419 ROIs. Our ROIs consisted of 400 cortical areas from the Schaefer parcellation defined in the HCP’s fs_LR space (Figure4A; Schaefer et al., 2018), and 19 subcortical regions defined in subject-level volumetric space using Freesurfer’s structural segmentation algorithm (Figure4B; Fischl, 2012). Regional BOLD signal fluctuation maps were first defined in each run as the standard deviation of resting-state BOLD signal averaged within each cortical or subcortical area of interest. The resulting matrices (N x 419) were averaged between runs within each session, which were then used to assess the relationship between time of day and regional BOLD signal fluctuation.

P-values were computed using 100,000 permutations, and corrected for multiple comparison using FDR-correction (q < 0.05). Results relating to cortical and subcortical ROIs were visualised in separate formats. Time of day effects on cortical regions were visualised on an fs_LR cortical surface mesh via Freeview, a part of the Freesurfer suite (Fischl, 2012). Each cortical region was assigned a colour-label (shown in Figure 4A) based on their correspondence to 17-large scale networks (Yeo et al., 2011; Schaefer et al., 2018) consisting of Temporo-Parietal, Default (A,B,C), Control (A,B,C), Limbic (A,B), Salience, Ventral Attention, Dorsal Attention (A,B), Somatomotor (A,B) and Visual (A,B). The results relating to the subcortical regions of interest were visualised in a heatmap. The 19 subcortical regions of interest consisted of left and right hemisphere masks of the nucleus accumbens, amygdala, hippocampus, caudate, thalamus, putamen, pallidum, ventral diencephalon, cerebellum and a single mask of the brain stem.

### 5.5 Time of day effects on resting state functional connectivity

We assessed the relationship between time of day and resting state functional connectivity (RSFC), and between RSFC and fluid intelligence. Fluid intelligence was chosen because it is a widely studied measure amongst resting-fMRI studies of brain-behavioural associations (Finn et al., 2015) and is one of the behavioural measures that is best predicted by resting-fMRI (He et al., 2018; Kong et al., 2019; Li et al., 2019). We computed a Pearson’s r correlation matrix (419 x 419), for each run of each subject using the mean time-series extracted from the 419 ROIs described above. The run-level correlation matrices were averaged within sessions to produce session-level estimates of RSFC for each subject. These matrices were then concatenated (N x 419 x 419) and correlated with time of day at each ROI to ROI pair (N x 419 x 419). We applied network-based statistics (Zalesky et al., 2010), a cluster-based thresholding technique, to control for family-wise error rate. We set an initial threshold of p < 0.001, and ran 100,000 permutations, while respecting family-structure of the participants (Winkler et al., 2015). P-values related to each correlation matrix underwent FDR-correction at q < 0.05. For all RSFC analyses, 5 subjects were excluded from Session 1 (N = 937 remaining) and 4 excluded from Session 2 (N = 865 remaining), due to missing fluid intelligence scores.

### 5.6 Global signal regression (GSR)

We were interested in the impact of GSR on our observations of time of day effects. To this end, we regressed the global signal from the ICA-FIX denoised CIFTI data. We defined global signal as the mean-time series computed from all cortical grayordinates (Kong et al., 2019; Li et al., 2019). We then repeated our analyses of the effects of time of day on regional BOLD signal fluctuation and on RSFC.

