## Supplementary Figures for "Time of day is associated with paradoxical reductions in global signal fluctuation and functional connectivity"

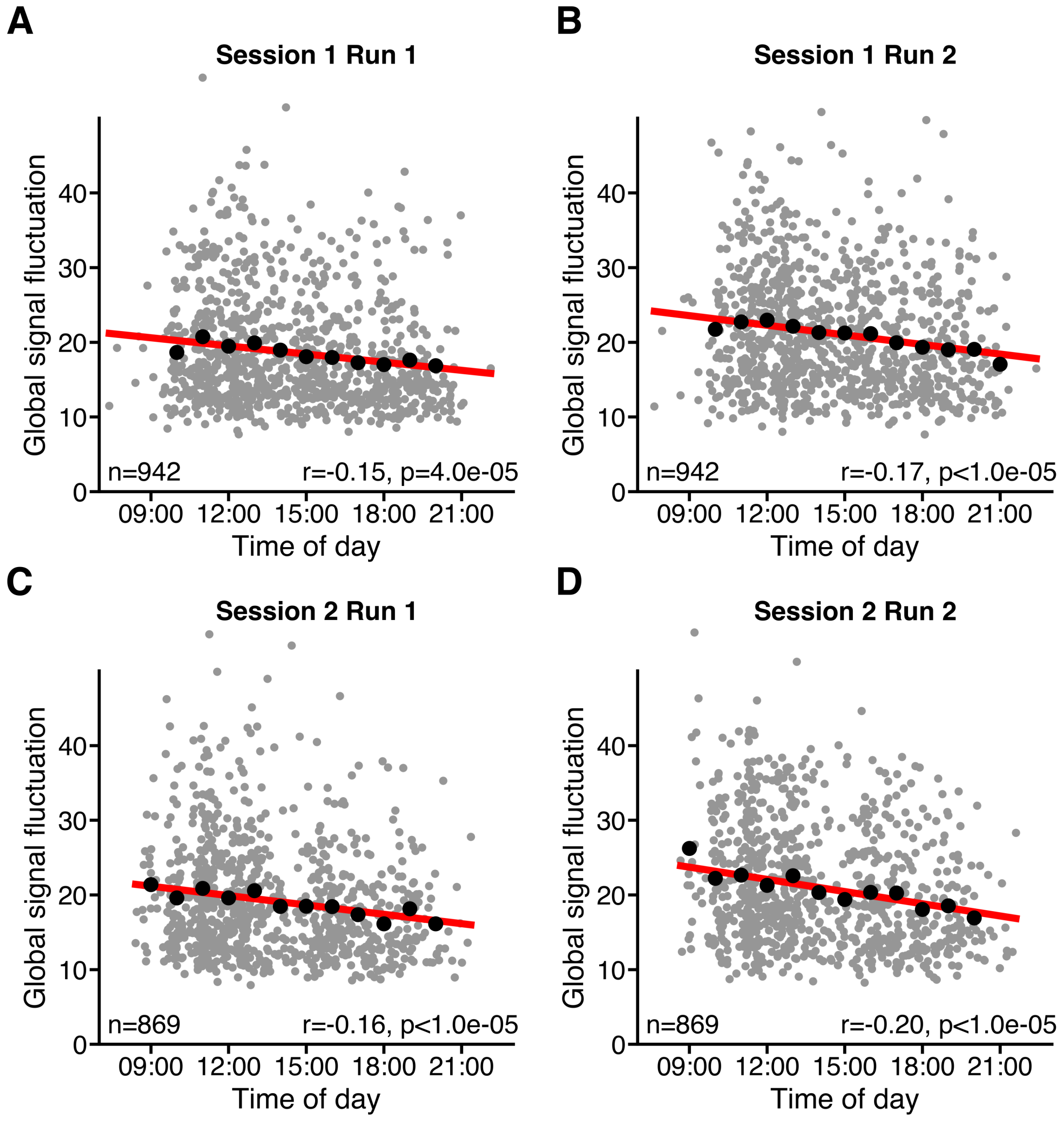


**Figure S1. Global signal fluctuation as a function of time of day shown separately for each run across two sessions. (A-D)** Grey dots denote individual subjects. Black dots show mean of GS fluctuation in hourly time windows. Line of best fit (red) was calculated based on data from all subjects in each plot. R values denote Pearson r correlation coefficient. P values were derived from 100,000 permutations while keeping family structure intact (Winkler et al., 2015). GS fluctuation was defined as the standard deviation of the global signal.

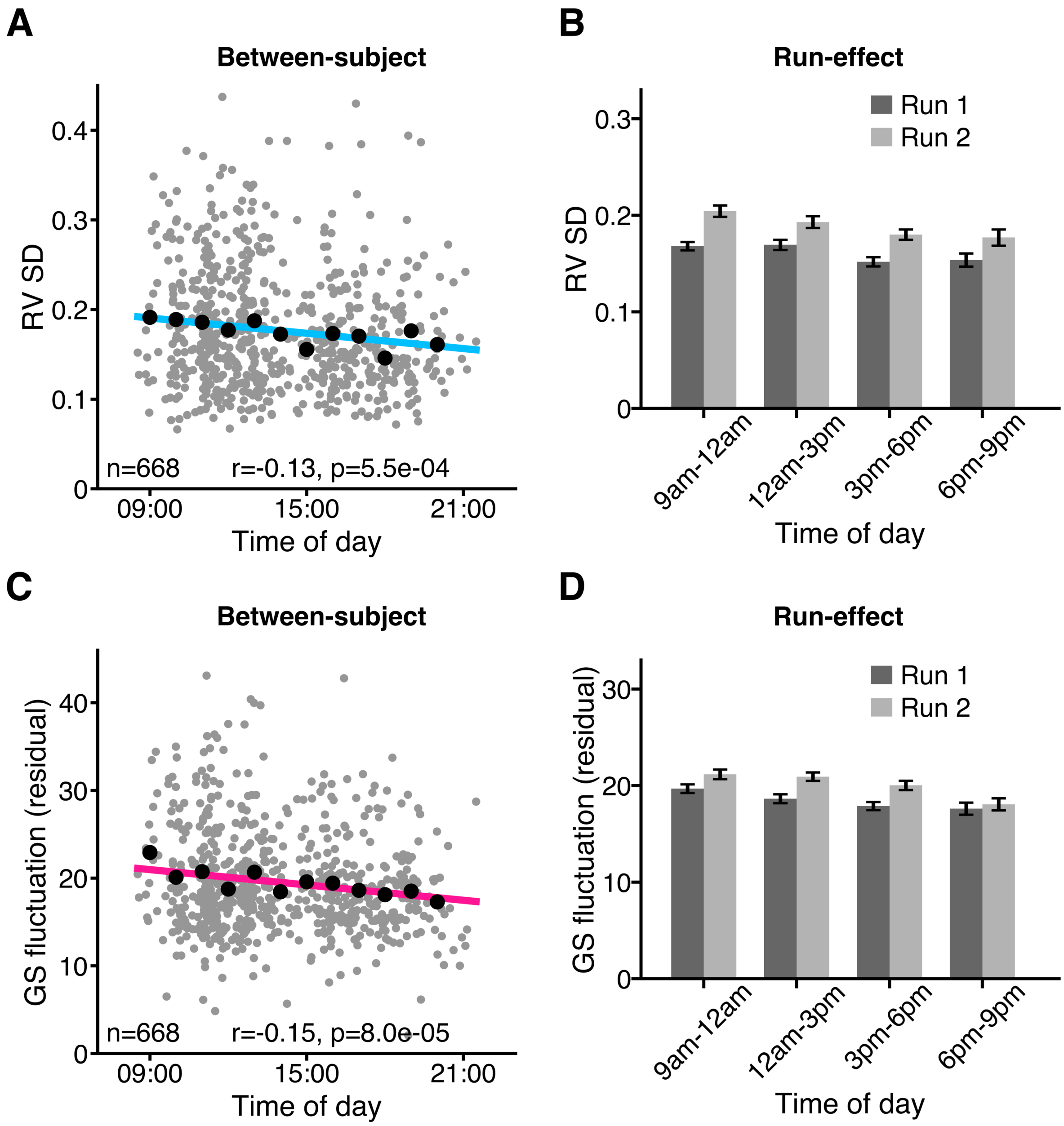


**Figure S2. Negative association between time of day and GS fluctuation remains significant after controlling for respiratory variation in Session 2. (A)** Between-subject variation of respiratory variation SD. **(B)** Run-effects on respiratory variation SD at different times of day. **(C)** Between-subject variation of GS fluctuation residual as a function of time of day. **(D)** Run-effects on GS fluctuation residual at different times of day. GS fluctuation residual was computed by group-level regression of respiratory variation SD from GS fluctuation. Grey dots denote individual subjects. Black dots denote mean of GS fluctuation in hourly (left), or 3-hourly time windows (centre). R values denote Pearson r correlation coefficients. P values were derived from 100,000 permutations while keeping family structure intact (Winkler et al., 2015). SD refers to standard deviation.

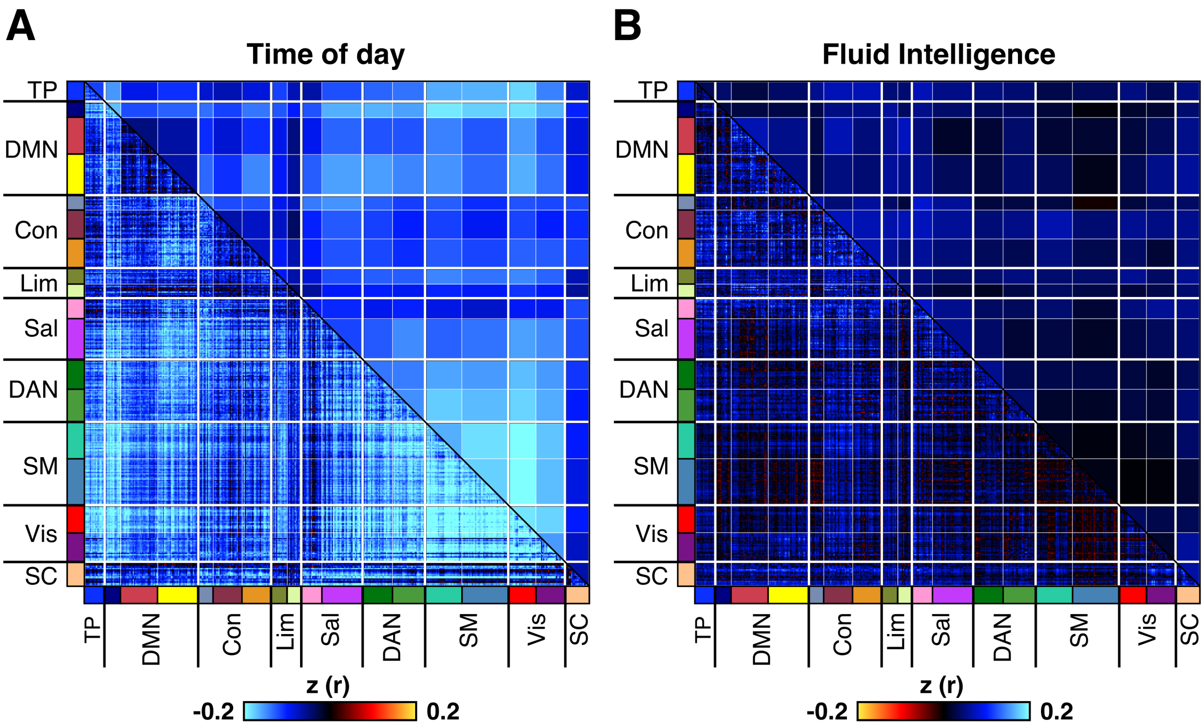


**Figure S3. Resting state functional connectivity (RSFC) region is negatively correlated with time of day across subjects in Session 2 (n = 865), with a magnitude that surpasses the strength of correlation between fluid intelligence and RSFC.** (A) Correlation between time of day and RSFC across subjects. (B) Correlation between fluid intelligence and RSFC across subjects. Colours in lower triangular of correlation matrix denote z-transformed Pearson r correlation coefficients. Colours in the upper triangular denote z-transformed r values from the lower triangular averaged within network pairs. Colours on label axes denote correspondence of 419 regions to 17 large-scale cortical networks and to subcortex (SC). Median absolute z values computed over the lower triangular were higher for time of day (0.13) than for fluid intelligence (0.04). Time of day - RSFC effects were significant, while RSFC - fluid intelligence effects were not significant for Session 2, as assessed by network-based statistics (FDR-corrected at $q < 0.05$). Note that the colourscale for the fluid intelligence - RSFC effects was inverted to facilitate visual comparison with time of day effects. For Session 1 results see Figure 6 in the main text.

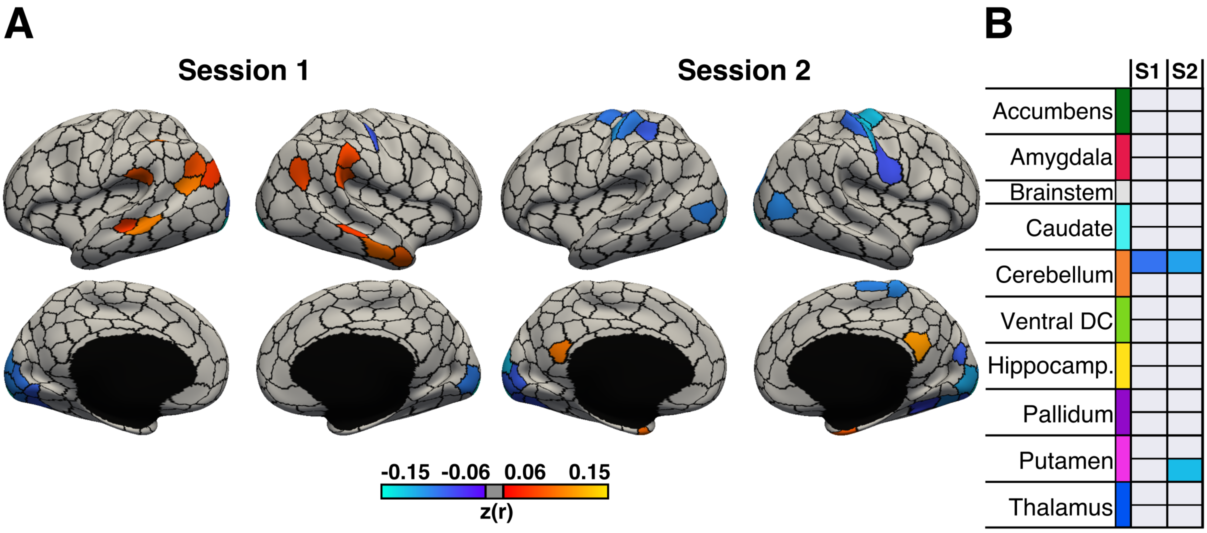


**Figure S4. Global signal regression results in an overall reduction of negative correlations between time of day and regional BOLD signal fluctuation across the brain, while introducing positive correlations at specific regions. (A)** Cortical regions showing significant correlations between time of day and BOLD signal fluctuation across subjects in Session 1 (n = 942) and Session 2 (n = 869). **(B)** Subcortical regions showing significant correlations time of day and BOLD signal fluctuation across subjects in Session 1 (S1) and Session 2 (S2). With the exception of brainstem, all subcortical regions are bilateral and presented as left-to-right hemisphere pairs (top to bottom). P values were derived from 100,000 permutations while keeping family structure intact (Winkler et al., 2015). Colours (cool-warm) denote cortical and subcortical brain regions with significant z transformed Pearson r coefficients ($q < 0.05$, FDR-corrected), while non-significant regions are shown in grey.

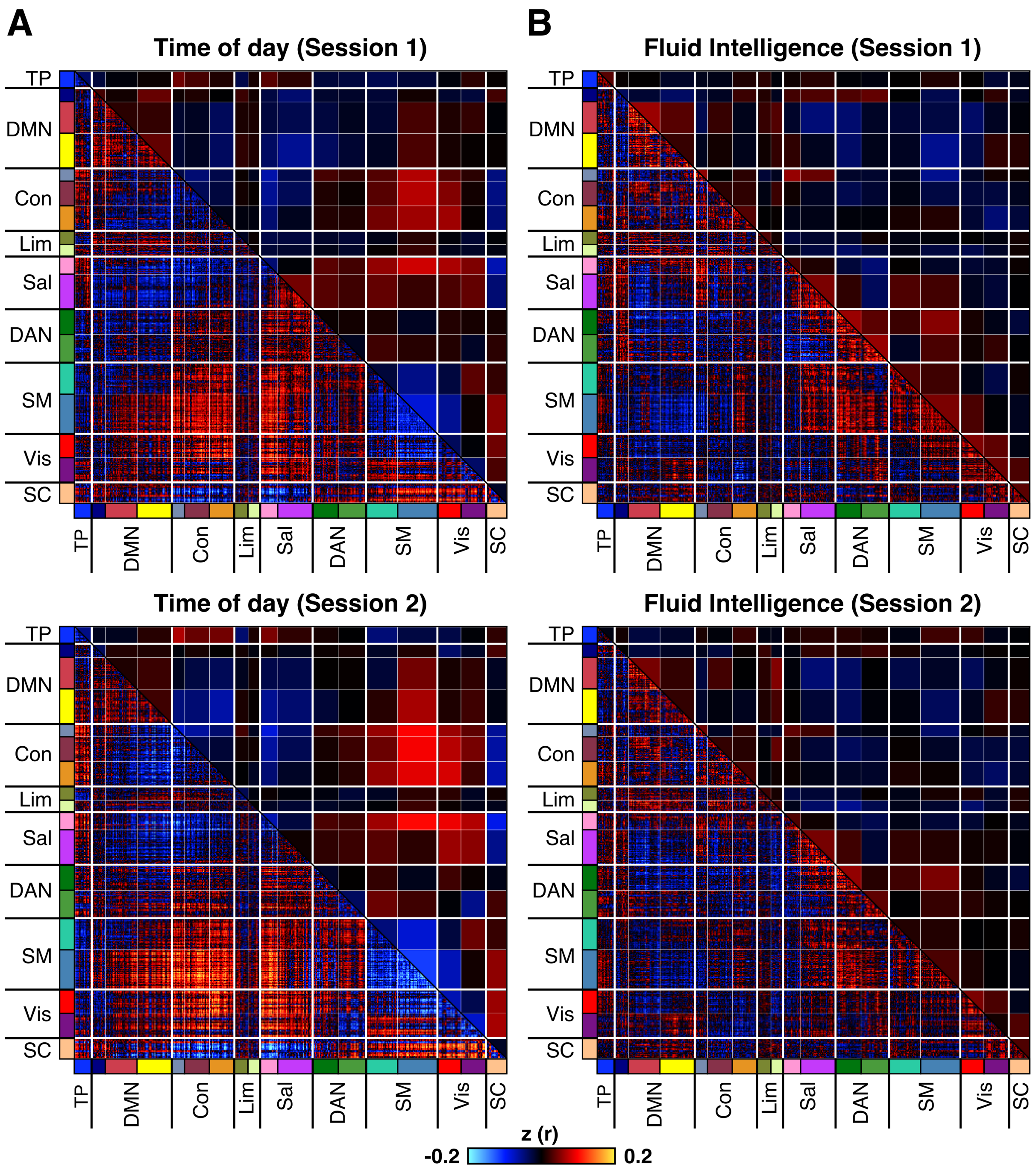


**Figure S5. Global signal regression reduces magnitude of negative correlations between RSFC and time of day, while introducing positive correlations for several large-scale circuits in both Session 1 (N = 937) and in Session 2 (N = 865).** (A) Correlation between time of day and RSFC across subjects. (B) Correlation between fluid intelligence and RSFC across subjects. Levels of correlation are visibly stronger between time of day and RSFC than between fluid intelligence and RSFC. Colours in lower triangular of correlation matrix denote z transformed Pearson r correlation coefficients. Colours in the upper triangular denote z values from the lower triangular averaged within network pairs. Colours on label axes denote correspondence of 419 regions to 17 large-scale cortical networks and to subcortex (SC). Time of day - RSFC effects were significant in both sessions as assessed by network-based statistics (FDR-corrected at $q < 0.05$), while fluid intelligence - RSFC effects were significant only in Session 1.

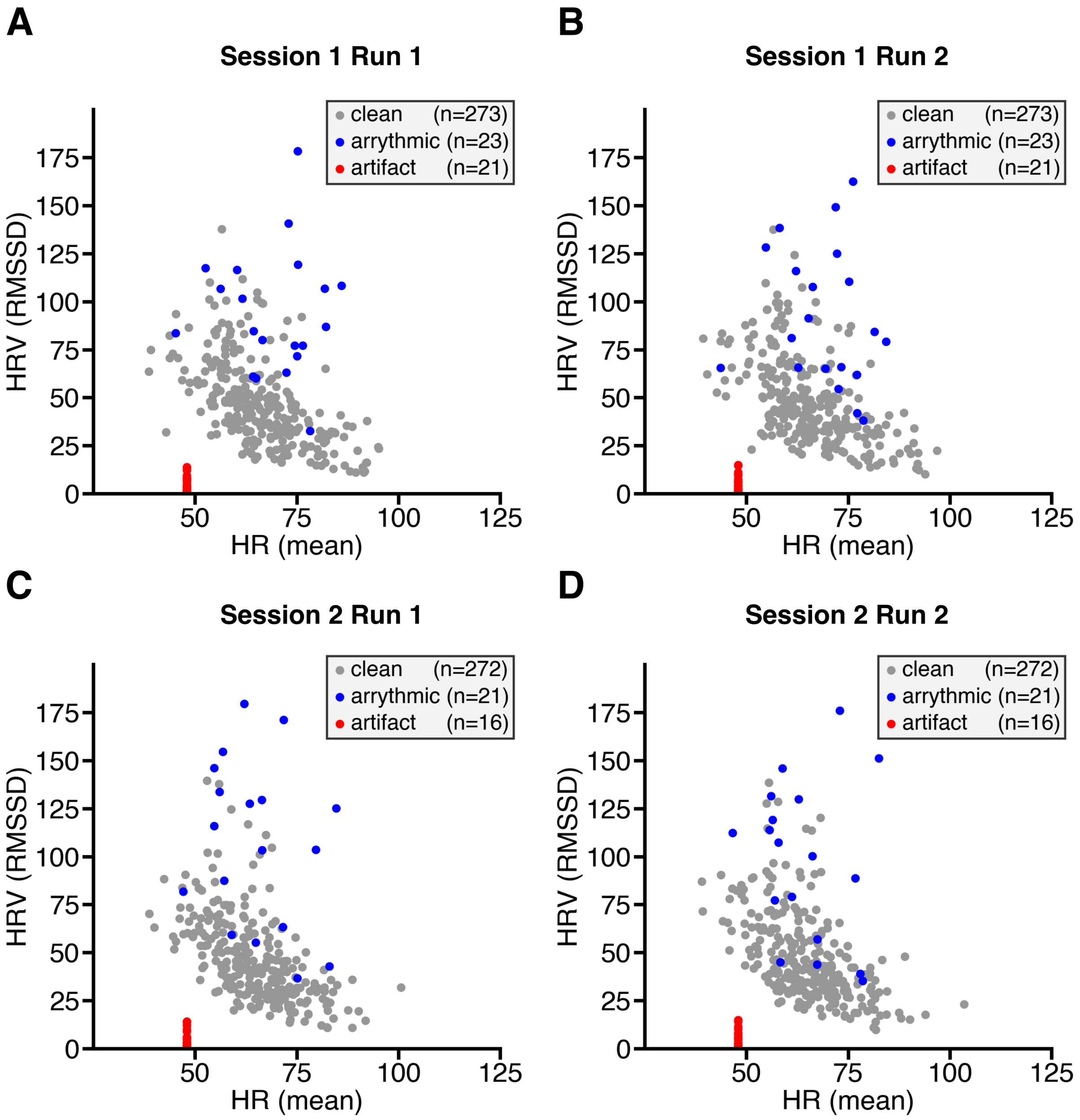


**Figure S6. Heart rate variability (HRV) as a function of Heart Rate (HR)**. Data is shown for subjects with pulse oximetry traces with sufficient quality to enable reliable peak detection. For a subject to be included in session-level analyses, both runs had to pass quality criteria. Despite good quality peak detection in these subjects, there were some additional anomalies. In several runs subjects exhibited a HR of exactly 48 beats per minute, with extreme low levels of heart rate variability, which were interpreted as likely artifactual in origin (shown in red). Other subjects (shown in blue) were identified as having arrhythmia of non-sinus origin based anomalous distributions on Poincaré plots (not shown). Some of these arrhythmic subjects are not visible on plot due to having extreme HRV (> 200 RMSSD). The remaining runs were deemed suitable for analysis of cardiac data (shown in grey). RMSSD - root mean square of the successive differences; HRV – heart rate variability.
